# Dynamic thermodynamic-informational entropic relationship (TIER) models of selective vulnerability to neurodegeneration

**DOI:** 10.64898/2026.04.08.714596

**Authors:** Peter S. Pressman, Cemal Basaran, Peter Foltz, Wan-Tai AuYeung, Joel Steele, Lisa Silbert, Lawrence E. Hunter

## Abstract

**BACKGROUND:** Neurodegenerative diseases share selective vulnerability patterns suggesting common physical mechanisms. We apply unified mechanics theory to neural systems, predicting that brain regions accumulate structural damage proportional to computational workload.

**METHODS:** We simulated a hierarchical neural network implementing relationships between mechanical work (W = F × D), proportional thermodynamic entropy accumulation (Δs ∝ W), and structural failure thresholds. Neural architectures at three hierarchical levels employed Hebbian learning across 2000 simulation sets, tracking thermodynamic entropy generation and dynamic stability. A coupled “siphon” model simulated cortical and subcortical support populations under constant cognitive demand.

**RESULTS:** Heteromodal integration nodes consistently exhibited elevated work, accelerated entropy accumulation, and dynamic instability across architectures. Support systems reached 50% population loss before cortical systems despite lower absolute work, demonstrating accelerated compensatory failure.

**DISCUSSION:** These thermodynamic-informational entropic relationship (TIER) models depict mechanisms underlying selective vulnerability across neurodegeneration, reframing neurodegeneration as the physical consequence of evolutionary trade-offs optimizing cognitive performance over longevity.

## INTRODUCTION

Neurodegenerative diseases have diverse anatomical pathways, histopathologies, prognoses and symptomatologies, but also share common properties that suggest common underlying principles. Age remains the strongest risk factor across neurodegeneration (1, 2). Pathology typically emerges in specific neural networks before spreading more broadly (3, 4). In hierarchically arranged cortices, atrophy is often most pronounced at poles or extremes. Cortex that has more recently evolved is often most prone to age-related atrophy (5, 6). Yet neurodegenerative conditions also share differences. Specific neural pathways are selectively vulnerable to specific proteinopathies, the reason for which is an active area of research (7, 8). We integrate insights from information theory, dynamic networks, unified mechanics theory, and behavioral neurology to investigate why such common properties emerge in complex biological information networks like the aging human brain.

We propose that these patterns emerge from the fundamental thermodynamic costs of information processing. All physical work, including the synaptic modifications required for information processing, carries risk of accumulated structural damage over time. According to the unified mechanics theory (9–17), the evolution of degradation in matter follows Boltzmann’s second law of thermodynamics (which Boltzmann called the second principle) (18). Under normal circumstances, systems do not fail from single catastrophic events but from accumulative irreversible thermodynamic entropy over their operational lifetimes (9–17). When applied to neural systems, unified mechanics theory predicts that brain regions that perform the most intensive information-related work will accumulate the most thermodynamic entropy, i.e damage, over time. This thermodynamic informational entropic relationship (TIER) reframes neurodegeneration as a systems-level failure to regulate thermodynamic entropy generation in the brain’s more computationally demanding regions.

Information processing in biological systems requires physical work at the cellular level. In neural networks, synaptic connections strengths represent the probability that one neuron will successfully influence another (19), embodying Shannon’s principle that information content inversely relates to probability (20). Learning often involves modifying these synaptic connections through Hebbian mechanisms (21), strengthening or weakening connections based on correlated activity. Each modification requires mechanical work, resulting in thermodynamic entropy generation (22), whether via receptor trafficking across membranes, dendritic spine formation and elimination, cytoskeletal reorganization, protein synthesis and transport, or other mechanisms (23–26). Every physical process contributing to synaptic modification carries its own thermodynamic cost, and these costs accumulate in proportion to work performed (Δs ∝ W) as a consequence of the second law applied to any system performing sustained work (9–17)

According to unified mechanics theory (12), materials fail when accumulated irreversible thermodynamic entropy reaches a critical threshold — the fracture fatigue entropy (FFE) — regardless of stress level, strain rate, or geometry (27, 28). FFE is not a single number but lies in a very narrow band, corresponding to a near 0.99 coordinate value of the Thermodynamic State Index (TSI), which tracks degradation on a continuum from zero at initial state to near one at failure; TSI cannot reach one because complete thermodynamic equilibrium is achieved only at infinity. Applied to neural systems, units processing the most informational work face the highest thermodynamic entropy generation rate, advancing most rapidly toward their FFE threshold.

We hypothesize that the brain’s convergent hierarchical organization amplifies these thermodynamic costs at higher hierarchical processing levels, especially when efferent axons traverse long distances and work is frequent. Mesulam’s sensory processing model describes information flow from primary sensory regions through unimodal association areas to heteromodal integration zones (29). Each hierarchical level must reconcile increasingly complex and diverse inputs, requiring synaptic modifications. Furthermore, heteromodal regions like the dorsal parietal cortex maintain some of the longest projection fibers in the brain, increasing the distance component of mechanical work as well (W = F × D) (30–32). The difference between metabolic expenditure and useful work is thermodynamic entropy, and TIER predicts that heteromodal integration hubs, with their long projection distances and sustained computational demands, will sit at the highest point on this mismatch curve. We hypothesize that the combination of intensive information integration and extensive physical connectivity creates disproportionate vulnerability to age-related failure, as thermodynamic efficiency inevitably declines over the system’s operational lifetime (12).

We further hypothesize that this vulnerability is extended to evolutionarily older structures through what we term the “siphon effect.” We model how, as primary cortical networks accumulate damage and lose efficiency, compensatory burden shifts to support systems, including the hippocampus, cholinergic nuclei, and noradrenergic projections from the locus coeruleus (33–36). These evolutionarily older structures possess limited volumetric reserve compared to distributed neocortex. Under increased workload from failing cortical systems, they accumulate thermodynamic entropy (disorder) rapidly, reaching fatigue failure entropy thresholds before the networks they support.

In any mechanical system, work inevitably generates irreversible entropy. An engine loses energy to friction. A bridge accumulates microfractures under repeated loading. Biological systems are no exception, but the sources of irreversibility are diverse. Rapid protein synthesis under demand produces misfolded byproducts requiring degradation. ATP hydrolysis powering molecular motors releases thermal energy that cannot be fully recaptured. Membrane recycling during vesicle trafficking is incomplete, accumulating lipid oxidation products. Cytoskeletal filaments sustain microdamage during the remodeling that physically reshapes synapses. No single mechanism is necessarily “the” cause of neuronal wear.

While FFE is not a single number, it lies in a very narrow band. FFE corresponds to a near 0.99 coordinate value of the Thermodynamic State Index (TSI) of unified mechanics theory. TSI values can only range between zero (at the initial state) and near one at the end. TSI cannot be one because a zero-entropy generation rate and complete thermodynamic equilibrium are achieved only in infinity. Systems fail when accumulated thermodynamic entropy reaches their intrinsic Fracture Fatigue Entropy band. When applied to neural systems, units that process the most informational work face the highest thermodynamic entropy generation rate.

## METHODS

### Theoretical Framework

#### Mechanical Work and Neural Failure

We apply the physical definition of mechanical work (W = F × D) to neural systems, in which force acts over distance during cellular processes. According to the second law of thermodynamics, no system can be 100% efficient. Total intake energy is used for intended good work, and the rest (the inefficiency part) is used for unintended work. The energy used for unintended work, (also referred to as energy unavailable for intended work) is the irreversible entropy that is responsible for damage. Therefore thermodynamic entropy (s) is related to work, (W): (Δs ∝ W), which in the brain relates to information processing. Synaptic modification repeatedly requires force over distance. This thermodynamic entropy is cumulative over time, leading to structural failure according to second law of thermodynamics.

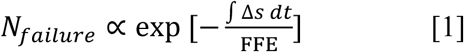

Equation 1 can be derived from the following Boltzmann’s second law of thermodynamics formulation, [9-14, 76]

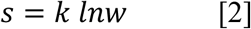

where *N_failure_* represents cycles to failure, Δ*s* is dissipated energy (thermodynamic entropy) per cycle, k is Boltzmann’s constant, *w* is the number of microstates reached and FFE (Fracture Fatigue Entropy) represents the critical entropy threshold at which failure occurs. FFE is an intrinsic material parameter representing the total entropy accumulation a system can tolerate before failure. When the accumulated entropy approaches FFE, the system approaches failure.

The unified mechanics theory has been extensively used across engineering systems from metals to particle filled polymer composites, concrete and electronics materials. It predicts that materials accumulate damage proportional to thermodynamic entropy until reaching failure. The ratio of accumulated entropy to FFE corresponds to the Thermodynamic State Index (TSI) of unified mechanics theory, which tracks degradation on a thermodynamic manifold from zero (initial state) to near one (failure).

#### Information Processing and Synaptic Modification

Neural information processing requires modifying synaptic weights, which represent connection probabilities between neurons. We implement a simplified Hebbian learning rule:

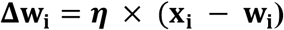

where **Δw**_**i**_represents synaptic weight change, **η** is the learning rate (0.1), **x**_**i**_ is the input (either external or from lower levels), and **w**_**i**_ is the current probability weight. This rule drives weights toward recent inputs, implementing unsupervised learning without pre-specified targets (37–39). While simpler than Oja’s rule (40), our approach achieves similar normalization through the hyperbolic tangent activation function, which bounds outputs to [-1, 1]. This prevents runaway weight growth while maintaining biological plausibility through activity-dependent synaptic modification.

Each weight modification requires mechanical work at the cellular level. We quantify this work as the cumulative magnitude of weight changes: W_node_ = Σ|Δw_i_|. This abstraction captures that synaptic modifications demand physical work without specifying the myriad cellular mechanisms involved. According to the unified mechanics theory, if the total entropy generation from all the myriad cellular mechanisms involved is calculated, their evolution would follow the second law of thermodynamics (equation 2), which underpins equation 1.

We use summation rather than vector operations because thermodynamic entropy is additive and locally generated. Each synaptic modification contributes irreversible entropy proportional to its magnitude regardless of coordinated activity in neighboring synapses. The dot product allows opposing weight changes to partially offset, which has no thermodynamic grounding. Entropy cannot cancel across sites by the second law. Squared weighting overemphasizes acute events, and rate of change captures instantaneous demand but loses cumulative history. Summation is therefore the only metric consistent with the second law’s requirement that irreversible entropy accumulate rather than cancel.

#### Connecting Information and Thermodynamics

Our models assume that information content relates to probability. Processing surprising inputs requires larger weight modifications. Processing many inputs per unit time requires more physical work than fewer inputs per unit time. Work requiring transportation of materials over long distances exceeds work required over short distances. The brain’s informational work reduces uncertainty at the cost of proportionate thermodynamic degradation.

The use of cumulative synaptic weight change as a proxy for mechanical, informational work has biological grounding. The actin cytoskeleton is the dominant structural element in dendritic spines. It occupies the spine head, undergoes continuous treadmilling, and its remodeling is necessary for lasting changes in synaptic strength (41, 42). With long term potentiation (LTP), spines enlarge and increase filamentous actin content. With long term depression (LTD), spines shrink and depolymerize actin. These structural changes track the sign and magnitude of weight modification (41). LTP induction shifts the equilibrium between globular and filamentous actin in proportion to the degree of spine enlargement(43), with the dynamic treadmilling pool providing the ATP-consuming mechanical substrate that executes these changes(44). Because actin polymerization is ATP-dependent and the degree of structural reorganization scales with the magnitude of the weight change required, thermodynamic work is proportional to |Δw_i_|.

While we focus on actin here, we recognize other mechanisms of plasticity such as AMPA receptor trafficking and postsynaptic density reorganization. The mechanistic details differ, but the thermodynamic principle does not. All candidate mechanisms require work, and thereby incur thermodynamic entropic risk. Analysis across five pre- and post-plasticity datasets confirms that plasticity updates energy budgets in proportion to |Δw|, with a precision-energy relationship of σ⁻² ∝ E⁵ (45). Motor proteins, including kinesin, dynein, and myosin, consume ATP proportional to distance traveled when delivering cargo to modification sites (46). Distal synapses therefore incur greater work for equivalent weight changes. The |Δw_i_| term in the model is a dimensionless index of molecular work per modification event, with the proportionality constant subsumed into the FFE threshold calibration. Future calibration studies linking measured ATP expenditure to specific |Δw_i_| magnitudes would allow quantitative translation of thermodynamic entropy units to biological energy equivalents.

Neurons carry substantial metabolic demands unrelated to information processing — ion pump restoration, protein turnover, membrane maintenance, and oxidative phosphorylation baseline. For the sake of model simplification, TIER treats these as approximately constant across cell types and hierarchical levels, focusing instead on the mechanical work of synaptic modification and axonal transport. TIER’s predictions derive entirely from this variable component. The framework applies to any physical system performing sustained work.

We track two distinct quantities. Informational entropy (H = -Σ p_i_ log(p_i_)) quantifies uncertainty in activation patterns.

### Computational Models

#### Hierarchical Network Architecture

We developed a simple convergent columnar architecture to model the thermodynamic consequences of convergent hierarchical architecture. The architecture distributes processing while maintaining hierarchical structure. Level 1 contains four primary nodes receiving independent inputs. Level 2 expands to four unimodal nodes providing redundant processing capacity. Level 3 contains four heteromodal nodes each receiving input from all unimodal nodes. This design models convergent hierarchical architecture, combining hierarchical position with systematic fan-in convergence at the highest level.

#### Network Implementation

All nodes use hyperbolic tangent activation: y = tanh(Σw_i_ × x_i_). We initialize weights uniformly on [0.1, 0.9] to avoid saturation.

We deliberately present random inputs drawn from [0.4, 0.6] with ±0.05 noise rather than structured signals. This design choice isolates the mechanical work inherent to hierarchical processing from confounds of stimulus or signal structure. By using random inputs, we demonstrate that work and therefore entropy accumulation emerges from network architecture alone, not from learning meaningful patterns.

This approach requires methodological clarification. In typical predictive networks learning structured signals, successful learning reduces uncertainty and informational entropy decreases over time. However, when learning random inputs, the network attempts to extract patterns from noise. This can increase uncertainty at higher hierarchical levels as the network integrates multiple random streams. How thermodynamic burden shifts under more structured, naturalistic inputs is addressed in a companion simulation using hierarchical melodic stimuli (195).

We track individual representative nodes from each level (not level aggregates) to reflect how work, and therefore thermodynamic entropy, accumulates in individual cells. Each simulation runs 20 learning iterations, repeated across 2000 independent trials with different random seeds.

Work quantification for each node: W(t) = Σ_i_ |w_i_(t) - w_i_(t-1)|, representing cumulative synaptic modifications. This per-node tracking captures that neurons fail individually based on their specific work history, not at the collective level. Stress, strain, and energy are defined at a material point rather than at the system level. Therefore, this definition is in agreement with the laws of unified mechanics theory. Regions with higher W(t) approach their FFE threshold sooner, as formalized in Equation 1.

#### Siphon Effect Model

The siphon model demonstrates how support systems fail before primary networks despite lower absolute work levels. We emphasize these simulations illustrate principles rather than quantitative biological predictions.

The system architecture consists of 1000 cortical cells representing a distributed processing network and 300 support cells representing focused nuclei such as the locus coeruleus. Individual capacity remains constant at C_per_cell_ = 1.0 across all cell types.

##### Work Dynamics

We impose constant cognitive demand (W_required_ = 3000 units) that must be maintained regardless of cell death. This prevents artifacts where declining function reduces work requirements.

Support systems provide baseline function plus compensation proportional to cortical dysfunction: W_support_ = W_baseline_ + 2.0 × (N_cortex_lost_/1000) × N_cortex_remaining,_ where W_baseline_ = 500, N_cortex_lost_ is the number of failed cortical cells, and N_cortex_remaining_ is the number of surviving cortical cells. This formulation creates peak support work at intermediate cortical loss, as support systems compensate for dysfunction while still having cortical cells to support. The factor of 2.0 represents the compensation intensity, scaling how much additional work support systems generate in response to cortical dysfunction, selected to clearly demonstrate the siphon effect principle within the simulation timeframe rather than to match specific biological parameters.

##### Failure Accumulation

Individual work intensity drives failure:

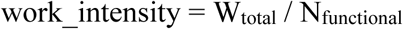

Accumulated work intensity integrates over time:

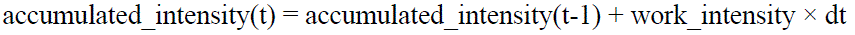

The theoretical basis for linking this accumulated work to system failure derives from the second law of thermodynamics. Thermodynamic entropy is additive and irreversible. Its evolution follows Boltzmann’s formulation:

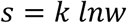

Where s is the total thermodynamic entropy. The number of microstates reached (disorder) as a function of total entropy, *s*, can be written as

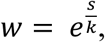

Because Δs ∝ W (Equation 1), accumulated work intensity corresponds directly to accumulated thermodynamic entropy. The scaling constant A absorbs the proportionality factor.

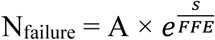

where A is a scaling constant to reflect baseline failure rate and N_failure_ represents the instantaneous failure rate. The following parameters are chosen to demonstrate a theoretical principle that systems with lower resilience fail before those they support under compensatory dynamics rather than to approximate specific biological values. We set FFE_cortex_ = 6.0 and FFE_support_ = 2.0, representing support systems’ lower resilience. These parameter values, including cell counts (1000 cortical, 300 support), work units (3000 required), compensation factor (2.0), and FFE ratios (6.0:2.0), are explicitly not calibrated to biological measurements or intended to approximate real neural populations, metabolic rates, or compensation magnitudes. Order-of-magnitude biological anchoring nonetheless supports the plausibility of these parameter relationships. The human locus coeruleus contains approximately 22,000–51,000 neurons bilaterally (47, 48), compared to approximately 10⁹–10¹⁰ cortical neurons it modulates, comprising a roughly 10⁴–10⁵ fold volumetric disparity corresponding to the C_per_cell ratio in our model. The FFE values assigned to cortical and support populations (FFE_cortex = 6.0; FFE_support = 2.0) are modeling parameters, not biological measurements. Their ratio reflects the qualitative observation that small, tonically active support nuclei with high per-neuron functional load and obligate oxidative byproducts of catecholamine synthesis face greater intrinsic damage accumulation than the distributed cortical populations they sustain. The sensitivity analysis confirms that failure order is invariant across all 30 tested FFE combinations, demonstrating that the siphon effect is a consequence of the differential itself rather than of any specific parameterization.

### Analysis Methods

For network models, we calculate three measures per node. Informational entropy quantifies uncertainty in activation patterns using Shannon entropy (20). Lyapunov exponents measure sensitivity to input perturbations with ε = 0.001 (49).

For siphon models, we track population dynamics over time, work intensity per surviving cell, accumulated work driving thermodynamic entropy generation, and system failure rates including critical transitions.

Statistical analyses use paired t-tests within architectures and independent t-tests between architectures. We report means ± standard deviations across 2000 simulation runs.

### Parameter Justification

Parameters were selected to demonstrate theoretical principles rather than match specific biological values. The learning rate (η = 0.1) allows convergence within 20 iterations. Network sizes are sufficient to show statistical patterns while remaining computationally tractable. The FFE ratio of 3:1 demonstrates differential vulnerability between systems. Time constants are scaled to show complete dynamics within the simulation window.

### Sensitivity Analysis

To assess the robustness of simulation results, we conducted a prespecified sensitivity analysis comprising six tests. First, work accumulation was quantified using four alternative metrics — Σ|Δw_i_| (primary), Σ|Δw_i_|², max|Δw_i_|, and d(Σ|Δw_i_|)/dt — applied in parallel to the columnar architecture. Second, structured inputs were tested using correlated Gaussian inputs (ρ = 0.3) and sinusoidal inputs (x_i_(t) = 0.5 + 0.4 × sin(2πf_i_t + φ_i_), distinct frequency per Level 1 node) as alternatives to the random input baseline. Third, FFE parameter space was swept across all 30 combinations of FFE_cortex values of 3.0, 4.0, 5.0, 6.0, 7.0, and 8.0 and FFE_support values of 1.0, 1.5, 2.0, 2.5, and 3.0, recording failure order and lead time at each combination. Fourth, compensation factor was varied across values of 0.5, 1.0, 1.5, 2.0, 2.5, 3.0, and 4.0 with all other parameters held constant. Fifth, activation function robustness was assessed by substituting tanh with sigmoid, ReLU, and linear activations. Sixth, parameter interactions were assessed using a factorial design across FFE ratio values of 2.0, 3.0, and 4.0, compensation factor values of 1.0, 2.0, and 3.0, and network sizes of 500, 1000, and 2000 cortical cells (27 combinations), with main effects and pairwise interaction plots computed for each parameter pair.

Code implementing all models is available at https://github.com/pspressman/TIER, including parameter files enabling reproduction and sensitivity analysis.

## RESULTS

Analysis across 2000 randomized input sets revealed distinct thermodynamic patterns across hierarchical levels. Across all hierarchical levels, thermodynamic entropy generation declined with learning as networks stabilized per sequential exposure. The magnitude of initial burden and the cumulative cost of achieving stability differed across levels, with Level 3 bearing the greatest thermodynamic cost of integration over repeated learning trials.

Level 3 nodes showed initial Lyapunov values of 0.13, compared to 0.05 for Level 1 and 0.07 for Level 2. All levels decreased over 20 iterations, with Level 3 showing the steepest decline (AUC: 0.55, SD: 0.16). Final values formed a V-shaped pattern: Level 1 highest (0.0044 ± 0.0002), Level 2 lowest (0.0022 ± 0.0001), and Level 3 intermediate (0.0034 ± 0.0001). All comparisons were significant (p < 0.001), with the strongest contrast between Levels 1-2 (t = 24.15).

Thermodynamic entropy generation in relation to mechanical work reflected this architectural difference. While Levels 1-2 matched the original model, Level 3 showed substantially higher initial energy demands (0.94 vs. 0.46) with nearly doubled area under curve (7.38 vs. 4.11), indicating concentrated early integrative work. After 2000 iterations, entropy curves showed tighter confidence intervals, demonstrating greater reliability in this distributed architecture (Figure 1).

**Figure 1:**
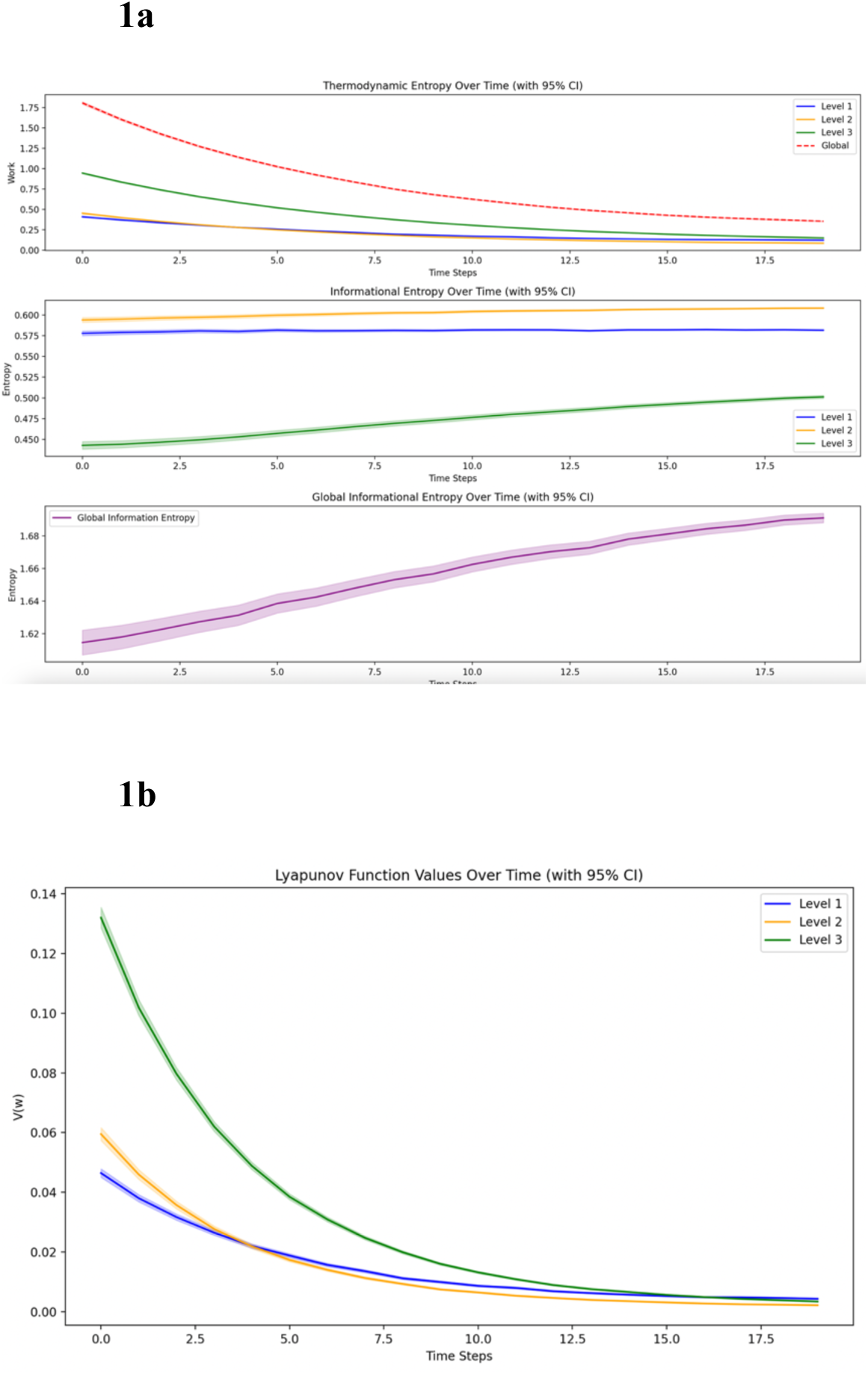
Columnar Network Dynamics. **1A.** Columnar network simulation across 2000 sets of 20 iterations. Thermodynamic entropy generation rate shows a sharp initial peak at Level 3, reflecting the front-loaded work of parallel integration. Though this work decreases with learning, Level 3 maintains a higher cumulative thermodynamic burden. **1B.** Lyapunov stability in the columnar network across 2000 sets of 20 iterations. The V-shaped pattern (Level 1 > Level 3 > Level 2) demonstrates how architectural choices influence dynamic stability.

### Siphon Effect Model

The extended TIER simulation demonstrated differential failure patterns between cortical and support cell populations over 20 time units. Support systems exhibited accelerated population decline compared to cortical systems in both absolute and relative terms. Support cells reached 50% population loss at 5.5 time units, while cortical cells reached equivalent decline at 11.5 time units. Support cell populations approached complete depletion (90% loss) at 5.9 time units, preceding cortical system collapse by 6.4 time units.

Individual support cells generated higher work intensity (and corresponding thermodynamic entropy) than cortical cells during critical decline phases. At the time when support systems reached 50% decline (5.5 time units), support cells sustained 3.4 units of work per cell compared to 2.5 units per cortical cell. Both systems eventually reached peak work intensity of 5.0 units per cell (the physiological maximum), but support systems reached this limit at 5.8 time units while cortical systems reached maximum intensity at 11.7 time units.

System failure rates confirmed accelerating failure dynamics through natural feedback loops between cell death, increasing individual workload, and structural damage accumulation.

Support systems reached peak failure rates of 538.1 cells per time unit at 5.9 time units, while cortical systems reached higher peak failure rates of 754.0 cells per time unit but much later at 12.5 time units. Support systems consistently reached failure thresholds 6.0 time units before cortical systems despite serving compensatory rather than primary processing roles (Figure 2).

**Figure 2:**
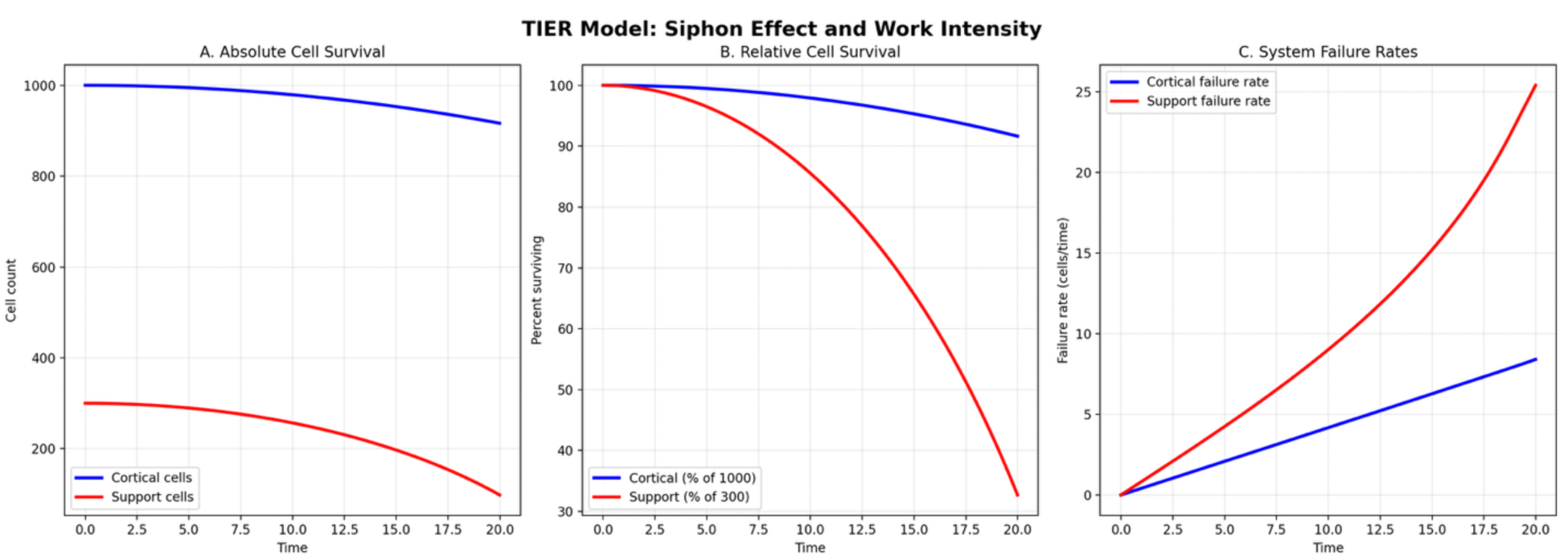
Siphon Effect Simulation. Three-panel plot modeling differential failure dynamics between cortical and supportive cell populations under constant cognitive demand. Absolute time values should be interpreted as illustrative of relative trajectories rather than biological time scales. **Panel A** (Absolute Cell Survival): Raw cell counts over time for cortical (blue) and support (red) populations. Support cells reach critical depletion well before cortical systems, despite beginning with lower absolute numbers, illustrating the accelerated failure trajectory predicted by the siphon model. **Panel B** (Relative Cell Survival): Survival expressed as percentage of initial population. Support systems reach 50% population loss substantially earlier than cortical systems, and approach complete depletion before cortical decline reaches equivalent levels. **Panel C** (System Failure Rates): Instantaneous failure rate per time unit for cortical and support populations. Support systems reach peak failure rates earlier and more steeply than cortical systems, reflecting the exponential acceleration of work intensity as population shrinks under constant demand.

### Sensitivity Analysis

The cumulative thermodynamic burden was robust across all work accumulation metrics. Level 3 exceeded Level 1 by 89.8% under Σ|Δw_i_| (p < 0.001), 254.3% under Σ|Δw_i_|² (p < 0.001), 101.2% under max|Δw_i_| (p < 0.001), and 86.3% under d(Σ|Δw_i_|)/dt (p < 0.001). The larger elevation under the squared metric is consistent with the theoretical expectation that overweighting large synaptic events amplifies the hierarchical gradient.

Under structured inputs, the gradient persisted under correlated Gaussian conditions (Level 3 elevation 102.9%, p < 0.001) and random input conditions (89.6%, p < 0.001). Sinusoidal inputs produced a gradient inversion of −9.4% (p < 0.001). The Oja learning rule implements online principal component analysis; a sinusoidal input has exactly one principal component by construction, and gradient inversion under maximally predictable input is the mathematically expected result. This finding is consistent with TIER’s prediction that structured, fully learnable inputs reduce thermodynamic burden: the sinusoidal condition represents the floor of that protection.

Support populations failed before cortical populations in all 30 of 30 FFE parameter combinations tested (FFE_cortex: 3.0–8.0; FFE_support: 1.0–3.0), with zero inversions. Lead time between support and cortical failure ranged from 0.3 to 10.7 time units, increasing monotonically with the FFE differential. Failure order was flat across the compensation factor range of 0.5–4.0, confirming that the siphon effect is driven by the FFE differential, not by coupling gain strength.

The hierarchical gradient held across all four activation functions, with Level 3 elevation ranging 11 percentage points across tanh, sigmoid, ReLU, and linear activations (all p < 0.001), confirming the result is a property of the convergent architecture rather than of activation function saturation.

Parameter interaction analysis identified FFE ratio as the dominant determinant of failure timing. Interaction plots showed near-parallel lines across all pairwise parameter combinations with no crossover interactions, indicating predominantly independent parameter effects and stable model behavior across the full parameter space.

## DISCUSSION

Our models illustrate costs to convergent hierarchical information processing. The network reflected a 105% increase in cumulative thermodynamic entropy generation at Level 3 compared to lower levels (AUC: 7.40 vs 3.61, p < 0.001). This elevation reflects how network topology shapes thermodynamic burden. Architectural features that amplify convergence amplify vulnerability.

With redundant access to all intermediate processing, heteromodal nodes correctly integrated information (informational entropy converging toward 0.50) while demanding nearly double the mechanical work of lower levels. Each node managing four direct inputs faced both increased structural burden and temporal cascade effects: nodes must reconcile four potentially conflicting representations of the same underlying information, generating additional thermodynamic entropy despite achieving lower informational uncertainty.

The Lyapunov stability analysis reveals that Level 3 nodes exhibited the highest initial instability and stabilized most rapidly over learning iterations. This differential stabilization reflects distinct computational constraints. Level 1 must respond to each new input. Levels 2 and 3 develop consistent output patterns regardless of input variability, but this stabilization requires continuous synaptic adjustment, corresponding to the elevated work and associated thermodynamic entropy we observed. Level 3 nodes achieve dynamic stability through sustained mechanical work rather than passive convergence. Heteromodal cortical regions, in other words, maintain high informational work demands even during routine processing because forcing coherent representations from variable inputs demands continuous synaptic modification.

### Biological Extensions

While these models primarily demonstrate the thermodynamic cost of convergent hierarchical architecture, biological heteromodal regions face additional workload from projection distance. Parietal to frontal connections are among the longest in the brain (31, 32). Connections spanning 10-15 centimeters perform orders of magnitude more work (W = F × D) than millimeter-scale primary projections. The disproportionate vulnerability of long-distance connectivity extends beyond the direct work of information transport. Long-distance fibers, despite comprising approximately 30% of neural connections, occupy 50 to 60% of brain volume, creating larger targets for diffuse injury while simultaneously requiring precise temporal synchronization, oscillatory coherence, and myelination integrity for effective function (50). Evolutionarily, both the hierarchy and distance relate to the same neocortical expansion that distinguishes humans from many other animal species but also establishes unique vulnerabilities.

This vulnerability can also be expressed in terms of mechanical systems’ complexity. In reliability engineering, the probability of system failure in series architectures grows exponentially with component count. A system of 1000 elements each 99% reliable retains only 0.004% system-level reliability(51). Perrow’s Normal Accident Theory formalizes this as a property of any system combining interactive complexity with tight coupling (52), conditions that precisely describe hierarchical sensory integration. Ashby’s Law of Requisite Variety further requires that a system’s controller must match the complexity of what it controls (53). These frameworks position the heteromodal cortex’s relatively high risk of thermodynamic entropic damage accumulation as directly relating to the combinatorial complexity of its inputs.

Our models focused on sensory hierarchies because sensory processing generates continuous, high-variance inputs that drive sustained synaptic modification. While subcortical motor systems maintain continuous tonic activity for postural and autonomic regulation, these circuits operate outside the heteromodal convergence hierarchy and generate relatively stereotyped outputs with low synaptic modification demand. Human motor cortex has a condition of baseline inhibition via basal ganglia. In contrast, sensory streams continuously stimulate the nervous system, ensuring constant informational work accumulation in heteromodal sensory regions, amplified by hierarchical organization and long efferent tracts. Heteromodal sensory cortex, which in Mesulam’s organization also involves hippocampus due to functional, cytological, and developmental similarities (29), is uniquely vulnerable both in healthy aging and conditions such as Alzheimer’s disease (54, 55). Neocortex is not, however, necessarily where pathology will first become apparent in neurodegenerative disease. While prefrontal and dorsoparietal cortices can lose volume at high rates even in healthy aging, early histopathological signatures of Alzheimer’s disease are seen elsewhere (56), such as the locus coeruleus and cholinergic nuclei (35, 36, 48). Both the locus coeruleus and the cholinergic system instrumentally support cortical sensory information processing (57–62).

The siphon model demonstrates how support systems for information integration may fail before the primary networks they sustain. As detailed in Results, support systems reached failure thresholds approximately 6.0 time units before cortical systems despite serving compensatory rather than primary processing roles. The mechanism emerges from coupled dynamics. As cortical efficiency declines, support systems increase activity to maintain function. With smaller populations and lower fracture fatigue entropy thresholds (FFE), support structures fail earlier before the networks they support, accelerating primary system failure through loss of compensation and recursive pathological protein seeding. Within this framework, every neural population traces a TSI trajectory from zero toward one. TIER predicts that heteromodal integration hubs and their support systems follow the steepest trajectories, reaching critical TSI values decades before primary sensory regions.

Cholinergic and noradrenergic structures are smaller and less flexible than neocortex, and fail in relation to projection length, cell volume, and evolutionary age (Table 1). The locus coeruleus has some of the longest projections in the brain and relatively few cells, and fails first, followed by the cholinergic systems (36, 48, 63). Work-related strains on the hippocampus are also substantial, relaying information between dynamic experiential instances and long-term semantic memories. This requires multiple strategies of resilience, including impressive flexibility and direct replacement of hippocampal cells to maintain FFE capacity (64–66). Larger and more dynamic than the cholinergic and noradrenergic systems (57, 67), the hippocampus is predicted to benefit from the siphon effect before itself falling victim to it.

**Table 1.**
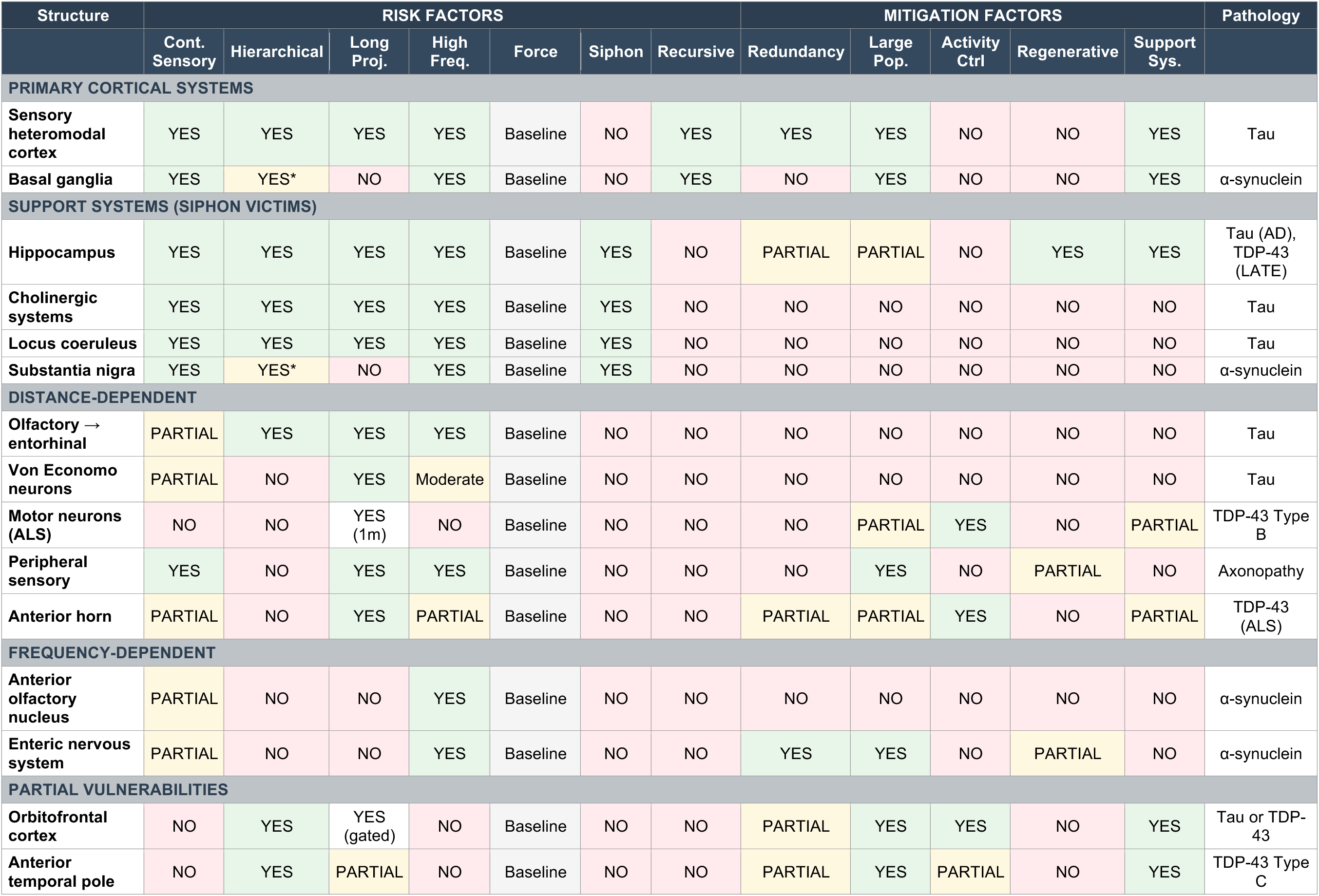
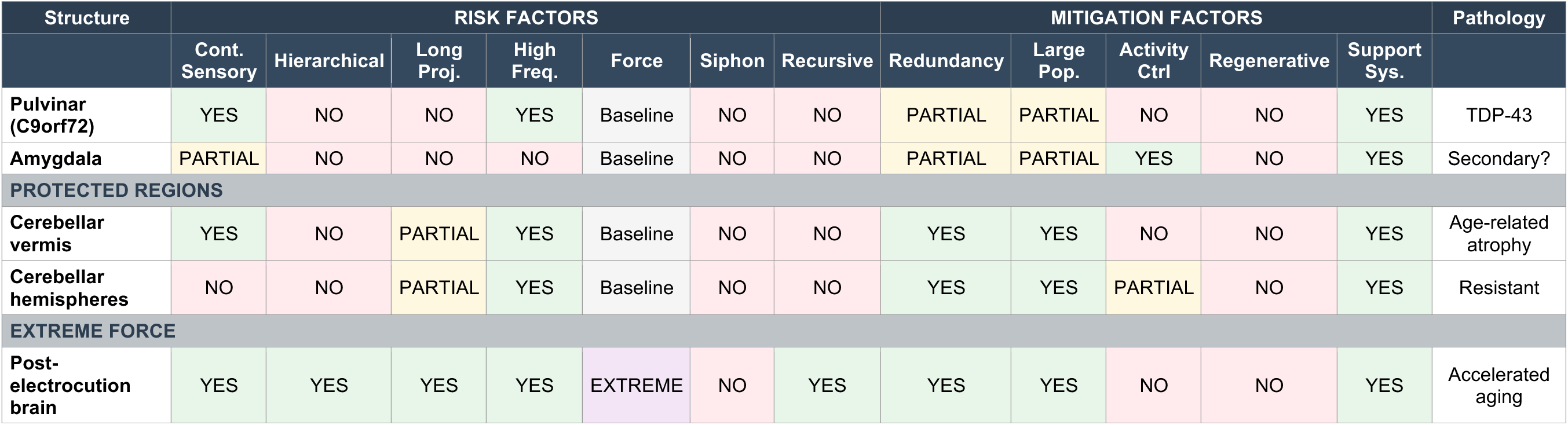
TIER Risk-Mitigation Matrix for Neural Structures. This table maps risk factors and protective mechanisms across neural structures according to the Thermodynamic-Informational Entropic Relationships (TIER) framework. Risk factors represent components of mechanical work accumulation (W = F × D over time), while mitigation factors represent fracture fatigue energy (FFE) or system resilience. Risk factors include: continuous sensory input (uninterrupted sensory processing that cannot be voluntarily suppressed); hierarchical processing (integration of information from multiple lower-level regions, with asterisks indicating inherited burden from cortical processing); long projections (axonal projections exceeding centimeter scale); high frequency (elevated synaptic cycling rates or continuous activity); force (mechanical force per operation, baseline for all structures except extreme cases); siphon burden (compensatory work inherited from failing primary systems); and recursive collapse (capacity to exhaust support systems that subsequently seed pathology back to the primary structure, enabling epidemic spread). Mitigation factors include: redundancy (parallel processing pathways versus bottleneck architecture); large population (substantial cell numbers providing reserve capacity); activity control (ability to gate, inhibit, or pause activity); regenerative capacity (cellular replacement or synaptic reorganization ability); and support systems (access to cholinergic, noradrenergic, or other modulatory support). Primary pathologies listed include tau, α-synuclein, TDP-43 (TAR DNA-binding protein 43) variants, and mixed pathologies. This simplified binary scoring (YES/NO/PARTIAL) represents a critical limitation, as it does not capture the relative weights of different factors. Recursive collapse likely confers orders of magnitude higher risk than individual vulnerability factors, while protective factors such as large cell populations may provide greater resilience than indicated by equal weighting. Future quantitative modeling incorporating disease prevalence data could establish appropriate weights for each factor. The table displays primary pathology only; all networked structures remain vulnerable to secondary pathological spread from connected regions. Abbreviations: AD, Alzheimer’s disease; ALS, amyotrophic lateral sclerosis; LATE, limbic-predominant age-related TDP-43 encephalopathy.

Our models are that of risk, not resilience, and so deliberately excluded protective mechanisms, including temporal buffering, recursive inhibition, sensory gating, predictive suppression, and more, to maintain focus on the fundamental threat demanding such an impressive defensive array (68–74). Beyond immediate survival benefits of energy efficiency, these myriad strategies are needed to prevent such complex integrative systems from rapidly accumulating overwhelming thermodynamic entropic burden.

### Metabolic convergence with TIER predictions

The difference between metabolic expenditure and useful work is thermodynamic entropy (Equation 2) (11, 18, 75–77). TIER predicts that heteromodal integration hubs, with their long projection distances and sustained computational demands, will sit at the highest point on this mismatch curve, and therefore will be the first regions where expenditure outpaces output in early disease, though they may then call on (siphon) supporting architecture.

While late Alzheimer’s disease is associated with hypometabolism as cell tissue has atrophied, in MCI and prodromal disease, the hippocampal formation demonstrates hypermetabolism negatively correlated with cognitive performance across multiple subdomains (78). Rubinski et al. (2020) extended these findings, showing that in MCI, higher regional tau-PET drives higher FDG-PET in frontal, temporal, and parietal association cortices, and that this hypermetabolism tracks with lower episodic memory (79). Despite consuming more fuel, the tissue is dysfunctional. Multiple independent mechanisms converge to widen this gap in the regions TIER predicts. Brain aerobic glycolysis, the glucose metabolism beyond oxidative needs that supports biosynthesis and synaptic remodeling, is highest in DMN hubs and declines with aging, eliminating repair capacity while computational demands persist (80–82). It is the loss of this channel, not total glucose or oxygen metabolism, that correlates with tau deposition in amyloid-positive individuals. The primary visual cortex illustrates the dissociation well, with high total energy consumption, low aerobic glycolysis, minimal amyloid, and minimal decline in typical Alzheimer’s. V1’s metabolic expenditure reflects high-throughput signal transmission through well-established circuits, not the continuous synaptic modification that drives TIER-relevant entropy generation.

On the supply side, multiple mechanisms compound the mismatch specifically in regions where TIER predicts the highest computational demand. Cerebral insulin resistance (“Type 3 diabetes,” (83, 84)) selectively starves the regions most dependent on continuous energy delivery. Neurovascular uncoupling disrupts the coupling mechanism itself: astrocytes mediate neurovascular signaling to capillary pericytes, controlling local blood flow in direct response to neural activity, but reactive astrogliosis disrupts this signaling so that energy demand persists or even increases while cerebral blood flow is reduced (85–87). Metabolic reprogramming through PKM2 replacing PKM1 shifts neurons to inefficient glycolysis, yielding less ATP per glucose molecule, more lactate, and more reactive oxygen species. Each of these mechanisms widens the gap between energy expenditure and useful work, disproportionately affecting high-demand regions TIER identifies as most vulnerable and compounding computational strain with metabolic inefficiency.

Recent evidence that myelin lipids serve as an active energy reserve during metabolic stress provides another consequential prediction of TIER. Prolonged metabolic stress creates substantial, reversible reductions in myelin water fraction across white matter (88), perhaps related to oligodendrocytes catabolizing myelin lipids through fatty acid β-oxidation to sustain axonal ATP and conduction when glucose is limiting (89). TIER predicts that this emergency metabolic pathway is chronically engaged in regions sustaining the highest cumulative informational workload. As local glucose supply fails to meet computational demand, myelin is consumed for energy, but the resulting demyelination increases the per-action-potential metabolic cost by orders of magnitude, accelerating glucose depletion and driving further myelin catabolism. This self-amplifying loop initiates in the late-myelinating association cortices where mechanical thermodynamic entropy generation is greatest and myelin reserves are thinnest. The APOE4 allele impairs cholesterol transport to oligodendrocyte membranes, disrupts astrocyte-derived lipid delivery essential for myelin maintenance, and blocks precursor cell differentiation (90, 91), crippling the repair arm of the cycle at multiple points simultaneously. APOE4 reduces the system’s capacity to reduce and recover from work performed over long distances, effectively lowering the FFE of the myelin infrastructure and shifting the threshold at which accumulated irreversible entropy produces failure.

Allostatic and bodily regulation demands contribute to the neural metabolic budget and cannot be entirely decoupled from information processing work. The present model treats these demands as approximately constant across anatomical regions, pending evidence of systematic regional variation that would generate differential vulnerability predictions independent of informational workload.

### Cellular and molecular extensions of TIER

The TIER framework extends into intracellular vulnerabilities through identical fundamental physics principles. Considering that work is a function of distance tau emerges as an especially vulnerable molecular structure in cells requiring long-distance information integration. Tau proteins maintain the structural integrity of microtubule highways under continuous mechanical loading from motor protein transport activity. While kinesin and dynein motor proteins perform direct work to move cellular cargo, tau bears cumulative irreversible entropy from stabilizing the transport infrastructure against these dynamic forces. Unlike motor proteins, tau is persistently engaged with microtubules, distributing multidirectional strain, including tension, shear, and torsion, across long distances and time periods. This continuous mechanical role, combined with tau’s high local concentration in long axons, its resistance to detachment under force, and its function as a damping element that absorbs bidirectional motor strain, suggests that tau may experience significantly more thermodynamic entropy generation per unit time than the transiently active motor proteins themselves (92–95). Furthermore, increasing evidence suggests that the initial vulnerability due to TIER principles also positions tau for a prion-like pathological cascade through neural networks, amplifying disease pathology through a process that is itself thermodynamic (96–100). Pathological protein propagation lowers FFE in connected nodes, accelerating their entropy accumulation and failure. As more vulnerable supportive structures struggle to compensate for cortical dysfunction, they accumulate pathological proteins at accelerated rates. When these overburdened support systems fail, they don’t simply withdraw their compensatory function but actively seed retrograde degeneration throughout the networks they were supporting.

These intracellular predictions map onto observed layer-specific vulnerabilities in the cortex. In entorhinal allocortex, where layer 4 is sparse, layer 2 reelin-positive neurons fail early due to continuous encoding demands and bottleneck architecture. Layer 3 pyramidal cells succumb from rapid bidirectional signaling requirements. Layer 5 neurons degenerate from extensive integration-distribution burden. Layers 1 and 6 remain relatively resilient through passive processing and modulatory rather than driving roles. In neocortex, layers 3 and 5 are especially involved with communication between parietal and frontal lobes, requiring long distances, and thereby also have predictably high levels of early Alzheimer’s pathology (101–105).

Amyloid, by contrast, is not as directly involved with the informational work of the cell and is predicted by TIER to be a diagnostic marker that, while toxic at pathological levels, represents a non-specific response to cellular stress rather than the primary driver of neurodegeneration.

Although amyloid presence is required for an Alzheimer’s diagnosis by definition (106), this diagnostic necessity may over-emphasize beta-amyloid’s true role in pathogenesis except in extraordinary cases. The most successful amyloid-clearing antibodies, lecanemab and donanemab, do demonstrate some clinical benefit and thereby support some toxicity of amyloid, yet their cognitive benefits remain remarkably modest despite convincing amyloid removal (107, 108). Meanwhile, 30-40% of cognitively normal elderly individuals harbor substantial amyloid deposits without symptoms, while tau pathology shows far stronger correlation with clinical progression (109).

Diverse cellular stressors, such as oxidative damage, inflammation, metabolic dysfunction, vascular insufficiency, infection, and trauma, all converge through stress-activated kinases JNK and p38 MAPK to upregulate BACE1, the rate-limiting enzyme for amyloid production (110). This non-specific response can occur rapidly, with amyloid plaques forming within hours of traumatic brain injury (111), and the peptide’s 400+ million years of evolutionary conservation and its protective functions at physiological concentrations support its origins as an ancient stress-response system that becomes toxic only when chronically overproduced (112).

Accordingly, neuroinflammatory markers such as microglial activation and astrocytic reactivity often precede amyloid deposition (113, 114), but like amyloid itself, these inflammatory responses may also represent non-specific markers of other underlying thermodynamic mechanisms. While only marginal benefits from amyloid removal are expected under TIER’s framework, in situations where amyloid is likely the primary driving force, e.g. rare genetic cases that increase amyloid production (115), TIER predicts that the benefit of anti-amyloid therapeutics will likely surpass those in the general population (116). These cellular, molecular, and metabolic extensions of TIER are detailed in Supplementary Table A.

### TIER and Selective Network Vulnerability Beyond Alzheimer’s Disease

The core formulations of TIER generate distinct predictions depending on which component dominates, and these predictions extend beyond Alzheimer’s disease to other forms of neurodegeneration as well. When thermodynamic entropy accumulates through synaptic frequency rather than projection distance, TIER predicts α-synuclein rather than tau as the vulnerable proteinopathy, as illustrated in Parkinson’s disease. Alpha synuclein perfectly embodies synaptic mechanical stress distinct from tau’s axonal burden. The protein regulates vesicle dynamics through repetitive nanometer scale conformational cycles. Each synaptic transmission event demands alpha synuclein bind curved vesicle membranes, facilitate SNARE complex assembly under mechanical tension, then dissociate for recycling (117–131). Basal ganglia perform rapid, continuous sensorimotor integration over short neural distances, and α-synuclein’s role in repetitive vesicle cycling creates irreversible entropy proportional to synaptic frequency (132–135). The olfactory system provides a natural experiment, with Alzheimer’s tau pathology emerging in long olfactory projections reaching entorhinal cortex while Parkinson’s α-synuclein concentrates millimeters from the bulb in the anterior olfactory nucleus, with similar findings in the gastrointestinal tract (117–130, 136–145). The same continuous sensory bombardment generates distinct molecular pathologies depending on whether mechanical work manifests as axonal transport stress or synaptic vesicle cycling. TDP-43 conditions, which each capture only partial TIER vulnerabilities — distance and sensory synapse in anterior temporal horn in ALS, hierarchy in semantic dementia’s predisposition for the anterior temporal lobe, sensory routing in *C9orf72*-related pulvinar thalamic atrophy— are correspondingly rarer, with the notable exception of limbic-predominant age-related TDP-43 encephalopathy (LATE), in which hippocampal convergence of multiple work drivers produces relatively high prevalence (146, 147). Frontotemporal dementia’s heterogeneity, its relative rarity compared to Alzheimer’s disease, and its preservation of subcortical cholinergic systems all follow from incomplete TIER risk profiles (e.g. convergence of sensory information on long distance Von Economo Neurons) and the absence of recursive cholinergic collapse (148–150).

TIER’s predictions extend beyond supratentorial dementia syndromes. The cerebellum may resist neurodegeneration despite extraordinary metabolic activity because parallel processing architecture and massive redundancy provide exceptional FFE capacity, though its vermis, receiving the most continuous vestibular sensory input, shows the highest vulnerability as predicted (151, 152) Motor neuron disease reflects projection length and synaptic integration risk even at spinal levels (153, 154), while the greater prevalence of sensory over motor peripheral neuropathy, and its increase with age, follow directly from the continuous nature of sensory work (155, 156). These and other predictions across proteinopathies, brain regions, and nervous system levels are mapped in Table 1 and detailed in Supplementary Tables B and C.

### Potential challenges to the TIER framework

Several apparent challenges to TIER resolve within the framework itself. If sensory integration work drives neurodegeneration, should sensory loss be protective? No, because having less information often means more, not less, cognitive work to establish understanding. In intact systems, diverse sensory inputs create stable ‘attractor states’ that enable efficient processing, as has been well-established across Hopfield networks (157–163). When a sensory modality degrades, this inhibition is lost. Remaining neurons must process noise and signal indiscriminately, multiplying mechanical work, because the brain’s drive to model reality completely persists despite degraded inputs.

Evidence that cognitive enrichment protects against dementia might appear to contradict TIER’s emphasis on work-driven vulnerability, but our simulations predict this effect directly.

Thermodynamic entropy generation decreases within each processing cycle as networks converge on efficient representations. Each successive exposure to similar information requires less synaptic modification as existing attractor states contextualize new inputs within established patterns. A cognitively enriched brain has built richer attractor landscapes, meaning that incoming information can be assimilated with less synaptic modification per unit of processing. Additionally, cognitive work builds synaptic redundancy and architectural flexibility, increasing FFE capacity. The columnar model especially illustrates both effects. Distributed connectivity produced lower informational entropy through more efficient computation, while the steepest decline in entropy generation occurred during early iterations. TIER predicts that this protection is greatest early in life, when both attractor formation and FFE capacity building are most efficient, and diminishes as cumulative thermodynamic entropy costs eventually overwhelm even well-maintained reserves, consistent with epidemiological evidence showing stronger protective effects of early-life cognitive enrichment than late-life intervention (164, 165).

Second, thermodynamic and informational entropy, while related, remain conceptually distinct. The relation between thermodynamic entropy and information entropy was studied in a seminal paper by Jaynes (166) where he stated that “The mere fact that the same mathematical expression occurs both in statistical mechanics and in information theory does not in itself establish any connection between these fields. This can be done only by finding new viewpoints from which thermodynamic entropy and information-theory entropy appear as the same concept.” He concludes that “The essential point in the arguments presented above is that we accept the von-Neumann —Shannon expression for entropy, very literally, as a measure of the amount of uncertainty represented by a probability distribution; thus entropy becomes the primitive concept with which we work [with], more fundamental even than energy.” TIER’s models represent the viewpoint Jaynes described. The brain’s hierarchical processing architecture creates a physical context in which thermodynamic entropy production and informational entropy reduction are causally linked through mechanical work at the synaptic level. The thermodynamic gradient is the price of reducing uncertainty.

TIER’s application of thermodynamic principles to neural systems invites the question of system boundaries. While thermodynamics typically speaks of closed systems, in practice, no physical system to which thermodynamics is usefully applied is perfectly closed. What matters is the degree of selective boundary control. The blood-brain barrier provides the brain a degree of thermodynamic isolation far exceeding most organs, but this isolation exists on a gradient. At specific anatomical sites, including the hippocampal vasculature, the olfactory epithelium, and the choroid plexus, the boundary weakens with age, increasing coupling to peripheral sources of entropy generation. Extravasated plasma proteins trigger inflammatory cascades, environmental particulates and pathogens access neural tissue directly through olfactory axons, and declining CSF turnover allows metabolic waste to accumulate. That said, relatively open boundaries alone seem insufficient to trigger neurodegeneration. Circumventricular organs such as the area postrema deliberately lack a blood-brain barrier yet are spared in neurodegeneration. The area postrema, however, performs simple chemosensory threshold detection with short local projections and minimal integrative demand. It may be that only where boundary degradation compounds an already-elevated thermodynamic burden — hippocampus (continuous memory encoding, high-frequency plasticity), olfactory cortex (constant environmental stimulation) — does the accelerated trajectory emerge. APOE4 may amplify this convergence by degrading BBB integrity via the cyclophilin A–MMP-9 pathway (increasing environmental entropy coupling)(167–170). The choroid plexus adds a further vicious cycle: age-related decline in CSF production coincides with increasing ventricular volume from brain atrophy, halving CSF turnover rate by the seventh decade. Waste products dwell longer in parenchyma even as the regions generating them are under greatest thermodynamic stress (171, 172).

Our models do not attempt to capture the myriad ways the brain may mitigate risk. Our goal here is to describe that risk, not mitigation. In a companion computational model implementing recursive descending inhibitory feedback across a seven-layer hierarchy trained on structured melodic stimuli, all three TIER measures increased with hierarchical level and no crossings at any learning step (173). A structural limitation of this network model also warrants acknowledgment. Level 3 nodes receive input from all four Level 2 nodes, a fan-in designed to illustrate the regional convergence property of heteromodal cortex rather than to replicate cellular-level synapse counts. The model deliberately abstracts regional architecture rather than cellular connectivity. Heteromodal cortex is defined by its position as a convergence point of multiple processing streams, and high fan-in at Level 3 models that regional property. As a consequence, the disproportionate thermodynamic burden at Level 3 relative to Level 2 reflects both hierarchical position and architectural convergence simultaneously, and the simulation cannot fully decompose these contributions.

The brain’s most complex and recently evolved information processors face the highest physical deterioration risk. While it’s the focus of our model, all neural information processing requires mechanical work at molecular scales, whether through receptor trafficking, protein conformational changes, ion channel gating, or cytoskeletal rearrangements, with force measurements consistently ranging from 1 to 1000 piconewtons over nanometer to micrometer distances (174, 175). Landauer’s limit establishes a universal minimum energy cost of kTln(2) per bit of information processing regardless of implementation (176). Regional vulnerability patterns persist across mechanisms: immediate early genes mediating the mechanical work of plasticity show highest expression precisely where our simulations identify greatest work-related thermodynamic burden (177–180). Finally, objections favoring specific mechanisms such as vascular failure or ribosomal stress over TIER miss its level of application. TIER does not replace molecular or physiological mechanisms but explains why diverse pathological processes converge toward shared endpoints under physical constraints. Entropy generation in molecular and physiological mechanisms can be used to compute entropy generation quantitatively for the TIER model The governing thermodynamic principles remain the same regardless of which mechanisms are measured.

### Predictions of the TIER Framework

TIER predicts that neurodegeneration risk is highest when workload is magnified by constant stimulation, hierarchical integration, long efferent distances, large forces, or low FFE. We have illustrated how the framework predicts progression, proteinopathy specificity, and prevalence across neurodegenerative diseases in proportion to the extent these risk factors are present. Supplementary Tables D and E detail some specific predictions across risk factors and therapeutic approaches. Here we highlight several that are particularly testable or surprising.

Because thermodynamic entropy generation is highest for information integration over long distances, disruption of the brain’s mitigating strategies for long-distance workload should increase neurodegeneration risk. Synchronized oscillations (181–183), sleep (184–186), and myelination integrity (187–189) all serve this function, and disruption of each has been correlated with subsequent neurodegeneration. TIER further predicts that atrophy of the cognitive cerebellum, itself involved in long-distance sensorimotor integration, will associate with risk of supratentorial neurodegeneration (190, 191).

More surprising predictions also emerge. If sensory integration is the most demanding synaptic work driving neurodegeneration, people with longstanding sensory integration deficits may be more prone to later neurodegeneration, as is increasingly demonstrated (192). TIER supports lifelong low-grade inflammation as a risk factor if it increases the work of sensorimotor integration, particularly through interference with effective myelination. This explains why chronic NSAID use over years consistently associates with decreased Alzheimer’s risk (193), while acute clinical trials of NSAIDs are ineffective (194). Chronic use reduces a lifetime of slowly accumulated inflammatory workload that cannot be rapidly reversed once tau cascades are underway. Similarly, 40 Hz gamma entrainment should reduce integratory workload by boosting long-distance oscillatory coherence, but would be most effective early, before significant tau accumulation, and using multimodal stimulation to reach heteromodal cortices (195, 196). Sleep enhancement should show particular efficacy when targeted to slow-wave phases, optimizing both glymphatic clearance and synaptic downscaling. TIER also supports bolstering noradrenergic and cholinergic systems as potentially underappreciated neuroprotective strategies even before Alzheimer’s pathology begins. Further predictions are listed in Supplementary Tables.

## Conclusion

The TIER framework reveals neurodegeneration as thermodynamic physics made manifest in biology, consistent with unified mechanics theory’s demonstration that Newtonian and thermodynamic principles govern the degradation of all physical systems (9–12). The brain’s most sophisticated and recently evolved capabilities contain unique risks. This understanding transforms how we approach brain health across the lifespan. Rather than searching for single treatments after cascade failures begin, TIER points toward precision identification of individual inflection points where work becomes destructive, optimizing efficiency before compensation overwhelms support systems, and tailoring treatment to unique expressions of Alzheimer’s and related disorders. Future therapies must be choreographed across the lifespan and individualized to individual risks.

In sum, TIER recasts Alzheimer’s as a systems-level failure to regulate thermodynamic entropy generation in the brain’s most demanding regions. Multiple vulnerabilities compromise the brain’s capacity to sustain complex integrative processing. These systems were never evolutionarily optimized for maintenance beyond reproductive age. As accumulated entropy advances irreversibly toward critical thresholds, mechanisms of resilience falter, resulting in the loss of informational stability in a brain built for brilliance but not permanence.

## Acknowledgements

Funding was provided by NIA K23 AG063900. Models and early manuscript drafting were assisted by Anthropic’s Claude and OpenAI’s ChatGPT.

