## Supplementary Table for "Dynamic thermodynamic-informational entropic relationship (TIER) models of selective vulnerability to neurodegeneration"

### TIER Predictions & Extensions Tables

#### Table A: Proteinopathy Predictions

| Proteinopathy | Predicted Mechanical Basis | Predicted Vulnerable Systems | Supporting Evidence | Refs |
|---|---|---|---|---|
| **Tau** | Maintains structural integrity of microtubule highways under continuous mechanical loading. Persistently engaged, distributing multidirectional strain (tension, shear, torsion). Cumulative stress ∝ projection distance (D). | Cells requiring long-distance integration: entorhinal cortex, heteromodal cortex (layers 3 & 5), long olfactory projections (35–50 mm axons). | High local concentration in long axons; resistance to detachment under force; damping element absorbing bidirectional motor strain; prion-like cascade potential. | 92–100 |
| **Alpha-synuclein** | Regulates vesicle dynamics via repetitive nanometer-scale conformational cycles. Each transmission: bind curved membranes, facilitate SNARE assembly under tension, dissociate. Cumulative work ∝ synaptic frequency. Low FFE: lacks stable secondary structure. | High-frequency synaptic cycling over short distances: basal ganglia, anterior olfactory nucleus, enteric nervous system. | Membrane binding regions experience repeated curvature stress. LC paradox resolved by gut-origin hypothesis: enteric α-syn propagates retrogradely via vagal connections. | 117–124, 137–145 |
| **TDP-43** | Multiple mechanical functions: nuclear-cytoplasmic shuttling, RNA granule assembly/disassembly, translational regulation. Different stresses trigger distinct aggregation pathways → multiple subtypes. | Each subtype captures one W aspect: Type B (ALS) = long projection; Type C (svPPA) = hierarchical; C9orf72/pulvinar = sensory routing. Exception: LATE (hippocampal convergence → higher prevalence). | Partial vulnerabilities explain rarity vs. AD. TDP-43 initiates at dendrites, consistent with synaptic integration burden. | 146–150 |
| **Amyloid-β** | Not directly involved with informational work. Non-specific stress response (fever analogy). Diverse stressors converge through JNK/p38 MAPK to upregulate BACE1. | All stressed regions (non-specific). | Modest cognitive benefits despite convincing removal. 30–40% cognitively normal elderly harbor amyloid. Forms within hours of TBI. 400+ million years evolutionary conservation. | 106–116 |

*Table A. Proteinopathy predictions. Each proteinopathy is mapped to its predicted mechanical basis (W = F × D), vulnerable neuronal populations, and supporting evidence. Reference numbers correspond to the main manuscript. Abbreviations: FFE, fold-fatigue endurance; BACE1, beta-site amyloid precursor protein cleaving enzyme 1; JNK, c-Jun N-terminal kinase; MAPK, mitogen-activated protein kinase; SNARE, soluble N-ethylmaleimide-sensitive factor attachment protein receptor; LC, locus coeruleus; svPPA, semantic variant primary progressive aphasia; ALS, amyotrophic lateral sclerosis; LATE, limbic-predominant age-related TDP-43 encephalopathy; TBI, traumatic brain injury.*

#### Table B: Regional & Structural Vulnerability

| System / Structure | TIER Risk Factors | TIER Prediction | Evidence | Refs |
|---|---|---|---|---|
| **Entorhinal cortex (layer-specific)** | Continuous encoding; bottleneck architecture; bidirectional signaling; integration-distribution burden | L2 reelin+ neurons fail early. L3: rapid bidirectional signaling. L5: integration-distribution burden. L1 & L6: resilient. | Observed laminar vulnerability in early AD. | 101–105, 125 |
| **Neocortical layers 3 and 5** | Communication between parietal and frontal lobes requiring long distances | Predictably high early AD pathology. | Observed. | 101–105, 125 |
| **Heteromodal sensory cortex (incl. hippocampus)** | Continuous sensory processing; hierarchical integration; long efferent tracts (10–15 cm); 2^level source scaling | Uniquely vulnerable in aging and AD. Most intensive processing → most rapid damage. | Observed. | 25, 29, 32, 54, 55 |
| **Hippocampus (metabolic)** | Central to cortical processing; highest BBB permeability with aging; peak insulin receptor density | Hypermetabolism in MCI preceding hypometabolic collapse. FDG-PET hypermetabolism negatively correlated with cognition. Levetiracetam confirms excess activity is harmful. | Apostolova 2018; Bakker 2012; Montagne 2015. | 78, 79, S1* |
| **DMN hubs (metabolic)** | Highest brain aerobic glycolysis; AG declines with aging; hub metabolism predicts downstream neurodegeneration | AG loss eliminates repair capacity. AG-tau correlation localizes vulnerability to mismatch, not total metabolism. Hub hypermetabolism mediates downstream neurodegeneration. | Goyal 2023; Vlassenko 2010, 2018; Galli 2025. | 80–82, S3* |
| **Locus coeruleus** | Longest projections; few cells; supports cortical processing; oxidative stress vulnerability | Fails first among support structures via siphon effect. | LC pathology among earliest in AD. | 32–34, 47, 48, 57–59 |
| **Cholinergic nuclei** | Support cortical sensory processing; smaller and less flexible than neocortex | Fail after LC, before hippocampus, via siphon effect. | Observed early cholinergic involvement. | 35, 36, 57–63 |
| **Hippocampus (structural)** | Central to cortical processing; relays experiential instances to long-term memories | Benefits from siphon before falling victim. Impressive FFE flexibility including cell replacement. | Hippocampal pathology in AD progression. | 64–67 |
| **Basal ganglia (PD)** | Rapid subcortical sensorimotor integration; short distances; high-frequency vesicle cycling | Vulnerable through frequency (not distance). Predicts α-syn over tau. | PD as disorder of rapid sensorimotor integration. | 132–135 |
| **Anterior olfactory nucleus** | Direct environmental bombardment; high-frequency; short distances | α-syn from intense local cycling. Same bombardment as AD olfactory but distinct proteinopathy due to short D. | PD pathology concentrates here, mm from bulb. | 126–131 |

| System / Structure | TIER Risk Factors | TIER Prediction | Evidence | Refs |
|---|---|---|---|---|
| **LC (PD context)** | Long projections → should predict tau, but accumulates α-syn | Resolved by gut-origin hypothesis: enteric α-syn → retrograde vagal propagation. | Emerging gut-origin evidence. | 137–145 |
| **Frontal cortex (FTD)** | Heteromodal sensory info; long projections; BUT BG gating reduces efferent frequency | Protected vs sensory cortex. Incomplete TIER vulnerabilities → diverse syndromes, relative rarity. | FTD much less common than AD. | 135, 148–150 |
| **Von Economo neurons** | Large spindle-shaped; exceptional distances; sparse distribution limits redundancy | High work-to-FFE ratios. In tau: VENs primary. In TDP-43: VENs secondary spread. | VENs die first in bvFTD. | 32, 148–150 |
| **Anterior vs. posterior insula** | Anterior: visceral, approximate. Posterior: cutaneous touch, mm precision, ms timing. | Opposing selective vulnerabilities at opposing ends of same structure. | Observed opposing patterns. | — |
| **FTD subcortical sparing** | Lacks LC, cholinergic, SN depletion | Without siphon effect targets, proteinopathy must arise endogenously in robust cortical systems → rarity. | ACh structures intact in FTD. | — |
| **Atypical AD (PCA, lvPPA)** | Primary cortical pathology without recursive collapse; still parietal | Rarer (de novo seeds in resilient tissue). Younger onset (with age, universal typical AD pathway). | Higher dyslexia rate in lvPPA suggests compensatory W. | S23*–S26* |
| **Cerebellum** | High metabolic activity; BUT parallel processing (no hierarchical burden); massive redundancy (80% of neurons = exceptional FFE) | Remarkable resistance. When degeneration occurs: vermis (70% vestibular afferents) most vulnerable. | Vermis: 4.59% GM loss/decade, nearly 2× overall rate. | 151, 152, 190, 191 |
| **ALS / Motor neurons** | Upper: long projections. Lower: integrate multiple sensory inputs. | Long neurons increase W through D. Anterior horn: synaptic processing. | Spasticity + weakness combination. | 153 |
| **Peripheral sensory vs. motor neuropathy** | Sensory: each sensation requires work at great distance. Motor: not every impulse requires motoric work. | Sensory neuropathy >> motor, increasing with age. Predicted co-occurrence with AD. | Observed sensory >> motor; increases with age; increased in AD. | 153–156 |
| **Post-electrocution brain** | Extreme force (F) applied to brain | Long-term cognitive decline even after improvement. Electric shock contributes to lifetime W. | Early cognitive decline years after injury. | S27*–S29* |

*Table B. Regional and structural vulnerability predictions derived from projection distance (D), synaptic frequency, force-generating load, fold-fatigue endurance (FFE), and the siphon effect (compensatory entropy transfer to support structures). Abbreviations: AD, Alzheimer's disease; PD, Parkinson's disease; FTD, frontotemporal dementia; BBB, blood-brain barrier; DMN, default mode network; AG, aerobic glycolysis; VEN, von Economo neuron; GM, gray matter; FDG-PET, fluorodeoxyglucose positron emission tomography; MCI, mild cognitive impairment; PCA, posterior cortical atrophy; lvPPA, logopenic variant primary progressive aphasia; bvFTD, behavioral variant frontotemporal dementia; ACh, acetylcholine; SN, substantia nigra; BG, basal ganglia.*

#### Table C: Risk Factor & Mitigation Predictions

| Risk Factor / Mitigation | TIER Mechanism | Prediction | Evidence Status | Refs |
|---|---|---|---|---|
| **Disrupted oscillations / E-I imbalance** | Oscillations and E-I balance mitigate long-distance integration workload | Disruption increases neurodegeneration risk. | Correlated with AD risk. | 181–183 |
| **Sleep disturbances** | Glymphatic clearance AND synaptic downscaling optimization | Disruption increases risk. Glymphatic impairment precedes Aβ deposition. AQP4 depolarization is central defect. | Correlated. Positive feedback loop. | 184–186 |
| **Demyelination** | Myelination eases integration over distance | Disruption increases long-distance workload → risk. | Correlated with subsequent risk. | 187–189 |
| **Demyelination (inflammaging)** | Chronic neuroinflammation damages myelin in long-range fibers. Temporal synchronization fails → hubs do MORE work to extract signal from misaligned inputs. | Increases $\dot{H}_k$ at hub by forcing noisier, less coherent input. Entropy cost of every integration cycle goes up. | HSV-1, periodontal, post-COVID all damage myelin. | 88–91 |
| **BBB breakdown** | Pericyte degeneration → loss of neurovascular coupling. Fibrinogen extravasation neurotoxic + inflammatory cascades. | Hippocampus is ground zero. BBB breakdown precedes Aβ/tau. APOE-ε4 carriers show elevated permeability without amyloid. | Montagne 2015; sPDGFRβ correlates with BBB leakage. | 167–170, S5* |
| **Microglial dysfunction** | Priming → hyperreactive. Dystrophic microglia lose phagocytic function. Complement-mediated synaptic stripping increases workload on remaining connections. | Each lost synapse increases burden on surviving synapses. Some FDG-PET 'hypermetabolism' reflects microglial glucose consumption for non-useful inflammatory cascades. | Xiang 2021; age-related neuroprotective → neurotoxic transition. | S4*, S11* |
| **Neurovascular uncoupling** | Reactive astrogliosis disrupts astrocyte-pericyte signaling. CBF no longer tracks neural activity. | Supply-demand mismatch at vascular delivery level. Coupling breaks before upply or demand individually fails. | McConnell 2019; Stackhouse & Mishra 2021; Mishra 2024. | 85–87 |
| **Aerobic glycolysis decline** | AG supports biosynthesis/remodeling. Highest in DMN hubs. Declines with aging. | AG loss—not total metabolism—correlates with tau. V1: high metabolism, low AG, minimal decline. Mismatch is the marker. | Goyal 2023; Vlassenko 2010, 2018. | 80–82 |
| **Cerebral insulin resistance ("Type 3 diabetes")** | Selectively starves high-demand regions. BUT: restoring fuel to broken coupling/AG/PKM2 system = more fuel burned inefficiently. | Restoring fuel without restoring efficiency: transient benefit then accelerating mismatch. Testable: insulin sensitizers alone plateau or worsen. | de la Monte 2008; Kciuk 2024; Bakker 2012 (reducing activity helped). | 83, 84, S1* |
| **PKM2 metabolic reprogramming** | PKM2 replaces PKM1 → inefficient glycolysis → less ATP/glucose, more lactate, more ROS. | Each mechanism widens the same gap. | Waller 2025. | S2* |

| Risk Factor / Mitigation | TIER Mechanism | Prediction | Evidence Status | Refs |
|---|---|---|---|---|
| **Inflammasome cascades (NLRP3)** | IL-1β, IL-6, TNF-α interfere with synaptic transmission/LTP. Gasdermin-D/pyroptosis. Anti-inflammatory decline removes brakes. | Inflammaging changes HOW FAST vulnerable regions reach threshold, not WHERE. | Inflammaging literature. | S7* |
| **Glymphatic impairment** | Waste accumulation degrades function. Regions generating most work harmed most by clearance failure. | Positive feedback: poor sleep → reduced clearance → more pathology → worse sleep. Partially reversible. | Progressive decline from middle age. Precedes Aβ. | S8* |
| **Vascular/metabolic inflammation (SVD)** | Penetrating end-arteries without collateral. Chronic hypoperfusion. Cumulative microinfarcts. | Deep white matter/periventricular zones. Midlife cardiovascular risk → late-life inflammaging substrate. | WMH, lacunar infarcts, microhemorrhages. | S9* |
| **Choroid plexus aging** | CP acquires Type I IFN signature → inflammatory factors into CSF. Reduced CSF production. | Periventricular structures bathed in inflammatory CSF. | Epithelial flattening, fibrosis, calcification. | S10* |
| **Sensory loss (hearing, vision, touch)** | Degraded input → predictions can't engage → gate fails → full inference loop per stimulus. Vicious cycle: +498% cost increase (195). | Sensory loss INCREASES risk. Hearing aids RESTORE INPUT QUALITY, allowing predictions to re-engage; quantified: vicious cycle +498% (195). | (195); epidemiology; S1*. | 155–163, S1* |
| **TBI as entropy accelerator** | DAI degrades gate. Surviving neurons work harder through degraded network. | Self-sustaining cascade follows DAI pattern, not stereotyped disease network. Latency = slow chronic mismatch. | Epidemiological TBI–dementia link; (195). | (173) |
| **Cognitive activity (early life)** | Work builds synapses → redundancy/flexibility → lifetime FFE. Richer attractor landscapes = less work per unit processing. | Protective. Simulations predict directly: work decreases as networks converge on efficient representations. | Observed. | 164, 165 |
| **Cognitive activity (late life)** | Benefits decrease as entropic costs overwhelm reserves. | Beneficial across lifespan but magnitude greatest earlier when attractor formation most efficient. | Lifelong risk prevention studies. | 164, 165 |
| **Cocktail party difficulty** | Difficulty distinguishing signal from noise increases work of sensory processing. | Predictable AD risk factor. | Increasingly demonstrated. | 192 |
| **Smoldering subclinical inflammation** | Low-grade chronic inflammation increases integration work via myelination interference. Changes HOW FAST, not WHERE. | Risk factor even without established risk factors. Clinical thresholds too high to detect indolent processes slowly increasing entropy over decades. | Consistent with inflammaging; subclinical threshold prediction not yet tested. | S7*, S11* |

| Risk Factor / Mitigation | TIER Mechanism | Prediction | Evidence Status | Refs |
|---|---|---|---|---|
| **Recursive collapse (siphon effect)** | Heteromodal processing siphons entropy to evolutionary predecessors → subcortical failure → retrograde degeneration | Path of least resistance. Explains AD/PD predominance. Evolutionary trade-off. | Siphon simulations; Galli 2025. | S3* |
| **Sensory loss paradox** | Loss disrupts attractor states → loss of recursive feedback inhibition → indiscriminate processing. | Sensory loss increases risk despite work driving degeneration. | Hearing loss as dementia risk factor. | 157–163 |
| **Premorbid sensory integration inefficiency** | Longstanding difficulty integrating multisensory information = more mechanical work per unit of processing across lifetime. | Motion sickness proneness, cocktail party difficulty, other integration deficits as prospective risk factors independent of frank sensory loss. | Cocktail party increasingly demonstrated. Motion sickness less studied as prospective marker. | 192 |
| **"Cognitive cerebellum" damage** | Cerebellum involved with long-distance sensorimotor integration | Atrophy predicts supratentorial neurodegeneration risk. | Less explored. | 190, 191 |

*Table C. Risk factor and mitigation predictions. Factors increasing cumulative work (W) or decreasing FFE are predicted to increase neurodegeneration risk; factors reducing work or enhancing resilience are predicted to be protective. Multiple mechanisms converge on the same thermodynamic gap between metabolic expenditure and useful computational work. Abbreviations: E-I, excitatory-inhibitory; AQP4, aquaporin-4; NLRP3, NLR family pyrin domain containing 3; IL, interleukin; TNF, tumor necrosis factor; BBB, blood-brain barrier; SVD, small vessel disease; IFN, interferon; CSF, cerebrospinal fluid; DAI, diffuse axonal injury; PKM, pyruvate kinase muscle isozyme; ROS, reactive oxygen species; LTP, long-term potentiation; WMH, white matter hyperintensities; CONTEXT, Contextual Organization of Noisy Topographic Encoding with eXponential Tuning (computational model).*

#### Table D: Therapeutic Predictions

| Intervention | TIER Rationale | Prediction | Evidence Status | Refs |
|---|---|---|---|---|
| **Chronic anti-inflammatory (NSAID distinction)** | Chronic use reduces lifetime accumulated inflammatory workload. Cannot be rapidly reversed once tau cascades underway. | Chronic use → decreased risk (observed). Acute trials → ineffective (observed). Effective window closes with tau. Benefit correlates with duration, inversely with tau burden. The chronic-vs-acute distinction is a direct TIER prediction. | Epidemiological chronic: consistent. Clinical acute trials: consistently negative. | 193, 194 |
| **40 Hz gamma entrainment** | Reduces integratory workload by boosting long-distance oscillation | Most effective early. Best multimodal. Optimal frequency varies by individual. Coherence > power. | — | 195, 196 |
| **Sleep enhancement** | Glymphatic clearance AND synaptic downscaling | Particular efficacy with slow-wave targeting, early disease. | — | — |
| **Bolstering noradrenergic/cholinergic** | Support systems for cortical processing | Potentially underappreciated neuroprotection even before AD pathology. | — | 35, 36, 57–61 |
| **Cholinergic augmentation in TBI** | ACh modulates cortical gain → sharpens predictions through surviving connections. | Partial compensation for gate degradation. Diminishes with severe DAI. Dose-response with connectivity loss. | David 2025 multicenter donepezil: memory improvement in chronic TBI. | S6* |
| **Amyloid-clearing antibodies** | Reducing secondary toxicity while fundamental fatigue failure continues. 'Antipyretics for fever in sepsis.' | Marginal in general population. Significantly greater benefit in dominantly inherited AD. DIAN-TU gantenerumab OLE: ~50% dementia risk reduction in longest-treated presymptomatic carriers (statistically inconclusive due to small sample/OLE design). | Lecanemab/donanemab modest in sporadic AD. Bateman et al. Lancet Neurol 2025. | 107–109, 115, 116 |
| **Insulin sensitizers / glucose alone** | Restoring fuel without efficiency = more fuel burned inefficiently → accelerates entropy. | Transient benefit then plateau/worsening. Must combine with coupling restoration or workload reduction. | Bakker 2012: reducing activity improved cognition. | 83, 84, S1* |
| **Levetiracetam** | Target: stop wasteful burning. Reduce activity in hypermetabolic but inefficient hippocampus → reduce entropy gap. | Most effective in early/MCI (hypermetabolism present). Loses efficacy in hypometabolic collapse (scar, not mechanism). | Bakker 2012: improved aMCI cognition. | S1* |
| **Neurovascular coupling restoration** | Targeting astrocyte-pericyte coupling > supply alone, because gap widens at delivery level. | Should outperform simple fuel delivery. Measurable as restored CBF-neural activity correlation. | Mishra lab. Testable. | 85–87 |

| Intervention | TIER Rationale | Prediction | Evidence Status | Refs |
|---|---|---|---|---|
| **Hearing aid / sensory restoration** | Restoring input allows predictions to re-engage, gate works, burden drops. Not enrichment—WORKLOAD REDUCTION. | Reduces metabolic load across cortical hierarchy, not just auditory cortex. Benefit diminishes with connection loss. | Computational modeling: +26% cost reduction with restoration (195). | (173) |
| **Familiar stimuli in dementia care** | Familiar stimuli restore gating via strong priors through degraded connections. | Measurably reduce metabolic load. Not 'stimulation'—enabling coasting on prediction. | (195); clinical music therapy observations. | (173) |
| **Combined approach** | Single-mechanism interventions: modest effects (multiple mechanisms widen same gap). Optimal = multiple angles simultaneously. | Coupling restoration + workload reduction + clearance enhancement should show synergistic, not additive, effects. | Testable. No direct evidence. | — |

*Table D. Therapeutic predictions. Interventions are evaluated by their predicted effect on the metabolic expenditure–useful work gap. Interventions targeting the mismatch directly (coupling restoration, workload reduction) are predicted to outperform those targeting fuel supply alone. Abbreviations: NSAID, non-steroidal anti-inflammatory drug; ACh, acetylcholine; TBI, traumatic brain injury; DAI, diffuse axonal injury; OLE, open-label extension; DIAN-TU, Dominantly Inherited Alzheimer Network Trials Unit; aMCI, amnestic mild cognitive impairment; CBF, cerebral blood flow; CONTEXT, Contextual Organization of Noisy Topographic Encoding with eXponential Tuning.*

#### Table E: Cellular, Molecular, and Metabolic Extensions

| Level | TIER Principle Applied | Prediction | Evidence | Refs |
|---|---|---|---|---|
| **Cortical layers (entorhinal)** | Complex integration/bidirectional neurons accumulate more damage than stereotyped processors | L2 (encoding bottleneck) > L3 (bidirectional) > L5 (integration-distribution) > L1, L6. | Observed laminar vulnerability. | 101–105, 125 |
| **Cortical layers (neocortex)** | Cells at bottlenecks with limited redundancy bear greater burden | Layers 3 and 5 (parietal-frontal long-distance) show high early AD pathology. | Observed. | 101–105, 125 |
| **Tau (intracellular)** | Fatigue failure extends intracellularly via W=F·D | Tau vulnerable in long-distance integration cells: persistently engaged, multidirectional strain, high concentration, resists detachment. | Tau > amyloid correlation with clinical progression. | 92–100, 109 |
| **IEG expression (Fos, Arc, BDNF)** | Regional vulnerability patterns persist across plasticity mechanisms | Highest in PFC, hippocampus, heteromodal hubs. Arc: AMPA trafficking; Fos: transcriptional programs; BDNF: cytoskeletal restructuring. | Concentrated in vulnerable regions. | 177–180 |
| **YWHAG:NPTX2 ratios** | Proteins regulating synaptic function via mechanical processes | Predict cognitive decline at sites of greatest adaptive work. | Recent findings. | 180 |
| **Non-Hebbian plasticity** | All neural processing requires mechanical work regardless of mechanism | Receptor trafficking, conformational changes, ion gating, cytoskeletal rearrangements: 1–1000 pN over nm–μm. Landauer's limit: kTln(2)/bit. | Force measurements across mechanisms. | 174–176 |
| **FDG-PET temporal arc** | Gap between expenditure and useful work IS thermodynamic entropy, visible on PET | Hypermetabolism (MCI) → destruction → hypometabolism (dementia). Classic pattern is scar, not mechanism. PCC/precuneus hypermetabolism is earliest conversion predictor. | Apostolova 2018; Rubinski 2020. | 78, 79 |
| **Microglial glucose uptake** | FDG-PET 'hypermetabolism' partly reflects inflammatory microglial activation, not neuronal work | Also mismatch: fuel for non-useful processes that worsen energy crisis. Metabolic gap includes non-neuronal waste. | Xiang 2021. | S4* |
| **Hub metabolism as siphon proxy** | Hub regions burning fuel to compensate for failing partners | Hub metabolic exhaustion predicts downstream collapse. Siphon effect measured with PET. | Galli 2025. | S3* |
| **V1 as TIER control region** | High energy consumption, low AG, minimal amyloid, minimal decline | Dissociates raw metabolic rate from TIER vulnerability. V1 processes intensively but stereotypically (low $\dot{H}_k$). | Vlassenko 2010, 2018; Goyal 2023. | 80–82 |

| Level | TIER Principle Applied | Prediction | Evidence | Refs |
|---|---|---|---|---|
| **Astrocyte-pericyte signaling** | Reactive astrogliosis disrupts neurovascular coupling | Coupling breaks BEFORE supply or demand individually fails. McConnell 2019 Figure 1: TIER restated in vascular terms. | McConnell 2019; Stackhouse & Mishra 2021; Mishra 2024. | 85–87 |
| **Metabolic reprogramming (PKM2)** | PKM2 replaces PKM1 → inefficient glycolysis | Each glucose molecule does less useful work. Combined with AG decline, insulin resistance, neurovascular uncoupling: every mechanism widens the same gap. | Waller 2025. | S2* |
| **CONTEXT model energy proxy** | $E(w) = E_0 + Em \times \lvert dw/dt \rvert$. Weight update magnitude IS metabolic cost. | Healthy (baseline), Hearing loss (+26%), Lost connections (+370%), Vicious cycle (+498%). Healthy = flat/low per note; vicious = flat/HIGH. | (195) | (173) |

*Table E. Cellular, molecular, and metabolic extensions. The work-entropy relationship (W = F × D) applies across scales from protein mechanics to whole-brain metabolic imaging. Abbreviations: IEG, immediate early gene; AMPA, α-amino-3-hydroxy-5-methyl-4-isoxazolepropionic acid; BDNF, brain-derived neurotrophic factor; PFC, prefrontal cortex; PKM, pyruvate kinase muscle isozyme; FDG-PET, fluorodeoxyglucose positron emission tomography; PCC, posterior cingulate cortex; MCI, mild cognitive impairment; V1, primary visual cortex; AG, aerobic glycolysis; CONTEXT, Contextual Organization of Noisy Topographic Encoding with eXponential Tuning.*

**Supplementary References**

Asterisked references (S#*) appear only in the supplementary tables and are not cited in the main manuscript text. All other reference numbers correspond to the main manuscript.

**S1*–S11*: Original Table-Only Citations**

**S1*** Bakker A, Krauss GL, Albert MS, Speck CL, Jones LR, Stark CE, Yassa MA, Bassett SS, Shelton AL, Gallagher M. Reduction of hippocampal hyperactivity improves cognition in amnestic mild cognitive impairment. Neuron. 2012;74(3):467-474. doi:10.1016/j.neuron.2012.03.023. PMID: 22578498.

**S2*** Waller TJ, Collins CA, Dus M. Pyruvate kinase deficiency links metabolic perturbations to neurodegeneration and axonal protection. Mol Metab. 2025;98:102187. PMID: 40505722.

**S3*** Galli A, Inglese M, Presotto L, Malito R, Di X, Toschi N, Pilotto A, Padovani A, Tassorelli C, Perani D, Sala A, Caminiti SP. Glucose metabolism in hyper-connected regions predicts neurodegeneration and speed of conversion in AD. Eur J Nucl Med Mol Imaging. 2025;52(12):4639-4651. PMID: 40471318.

**S4*** Xiang X, Wind K, Wiedemann T, Blume T, Shi Y, Briel N, et al. Microglial activation states drive glucose uptake and FDG-PET alterations in neurodegenerative diseases. Sci Transl Med. 2021;13(615):eabe5640. PMID: 34644146.

**S5*** Montagne A, Barnes SR, Sweeney MD, Halliday MR, Sagare AP, Zhao Z, Toga AW, Jacobs RE, Liu CY, Amezcua L, Harrington MG, Chui HC, Law M, Zlokovic BV. Blood-brain barrier breakdown in the aging human hippocampus. Neuron. 2015;85(2):296-302. doi:10.1016/j.neuron.2014.12.032. PMID: 25611508.

**S6*** Arciniegas DB, Almeida EJ, Sander AM, Bogaards JA, Giacino JT, Hammond FM, Harrison-Felix CL, Hart T, Ketchum JM, Mellick DC, Sherer M, Whyte J, Zafonte RD. Multicenter Evaluation of Memory Remediation in Traumatic Brain Injury With Donepezil: A Randomized Controlled Trial. J Neuropsychiatry Clin Neurosci. 2025;37(2):102-114. doi:10.1176/appi.neuropsych.20230055. PMID: 39628282.

**S7*** Heneka MT, Kummer MP, Stutz A, Delekate A, Schwartz S, Vieira-Saecker A, Griep A, Axt D, Remus A, Tzeng TC, Gelpi E, Halle A, Korte M, Latz E, Golenbock DT. NLRP3 is activated in Alzheimer’s disease and contributes to pathology in APP/PS1 mice. Nature. 2013;493(7434):674-678. doi:10.1038/nature11729. PMID: 23254930.

**S8*** Iliff JJ, Wang M, Liao Y, Plogg BA, Peng W, Gundersen GA, Benveniste H, Vates GE, Deane R, Goldman SA, Nagelhus EA, Nedergaard M. A paravascular pathway facilitates CSF flow through the brain parenchyma and the clearance of interstitial solutes, including amyloid β. Sci Transl Med. 2012;4(147):147ra111. doi:10.1126/scitranslmed.3003748. PMID: 22896675.

**S9*** Senatorov VV Jr, Friedman AR, Milikovsky DZ, Ofer J, Saar-Ashkenazy R, Charbash A, Jahan N, Chin G, Mihaly E, Lin JM, Ramsay HJ, Moghbel A, Preininger MK, Eddings CR, Harrison HV, Patel R, Shen Y, Ghanim H, Sheng H, Veksler R, Sudmant PH, Becker A, Hart B, Rogawski MA, Dillin A, Bhatt DK, Kaufer D. Blood-brain barrier dysfunction in aging induces hyperactivation of TGFβ signaling and chronic yet reversible neural dysfunction. Sci Transl Med. 2019;11(502):eaaw8283. doi:10.1126/scitranslmed.aaw8283. PMID: 31366449.

**S10*** Baruch K, Deczkowska A, David E, Castellano JM, Miller O, Kertser A, Berkutzki T, Barnett-Itzhaki Z, Bezalel D, Wyss-Coray T, Amit I, Schwartz M. Aging-induced type I interferon response at the choroid plexus negatively affects brain function. Science. 2014;346(6205):89-93. doi:10.1126/science.1252945. PMID: 25147279.

**S11*** Perry VH, Holmes C. Microglial priming in neurodegenerative disease. Nat Rev Neurol. 2014;10(4):217-224. doi:10.1038/nrneurol.2014.38. PMID: 24638131.

**S12*–S18*: Migrated from Main Manuscript (Former Refs 142–148)**

**S12*** Beach TG, Adler CH, Sue LI, Shill HA, Driver-Dunckley E, Mehta SH, et al. Vagus nerve and stomach synucleinopathy in Parkinson's disease, incidental Lewy body disease, and normal elderly subjects: evidence against the "body-first" hypothesis. Journal of Parkinson's Disease. 2021;11(4):1833-1843.

**S13*** Borghammer P, Van Den Berge N. Brain-first versus gut-first Parkinson's disease: a hypothesis. Journal of Parkinson's Disease. 2019;9(S2):S281-S295.

**S14*** Bottner M, Zorenkov D, Hellwig I, Barrenschee M, Harde J, Fricke T, et al. Expression pattern and localization of alpha-synuclein in the human enteric nervous system. Neurobiology of Disease. 2012;48(3):474-480.

**S15*** Braak H, de Vos RAI, Bohl J, Del Tredici K. Gastric α-synuclein immunoreactive inclusions in Meissner's and Auerbach's plexuses in cases staged for Parkinson's disease-related brain pathology. Neuroscience Letters. 2006;396(1):67-72.

**S16*** Braak H, Rüb U, Gai WP, Del Tredici K. Idiopathic Parkinson's disease: possible routes by which vulnerable neuronal types may be subject to neuroinvasion by an unknown pathogen. Journal of Neural Transmission. 2003;110(5):517-536.

**S17*** Hilton D, Stephens M, Kirk L, Edwards P, Potter R, Zajicek J, et al. Accumulation of α-synuclein in the bowel of patients in the pre-clinical phase of Parkinson's disease. Acta Neuropathologica. 2014;127(2):235-241.

**S18*** Holmqvist S, Chutna O, Bousset L, Aldrin-Kirk P, Li W, Björklund T, et al. Direct evidence of Parkinson pathology spread from the gastrointestinal tract to the brain in rats. Acta Neuropathologica. 2014;128(6):805-820.

**S19*–S22*: Sensory Integration and Inflammaging**

**S19*** Deal JA, Betz J, Yaffe K, Harris T, Purchase-Helzner E, Satterfield S, Pratt S, Govil N, Simonsick EM, Lin FR; Health ABC Study Group. Hearing Impairment and Incident Dementia and Cognitive Decline in Older Adults: The Health ABC Study. J Gerontol A Biol Sci Med Sci. 2017;72(5):703-709. doi:10.1093/gerona/glw069. PMID: 27071780.

**S20*** Lin FR, Albert M. Hearing loss and dementia — who is listening? Aging Ment Health. 2014;18(6):671-673. doi:10.1080/13607863.2014.915924. PMID: 24875093.

**S21*** Brenowitz WD, Kaup AR, Lin FR, Yaffe K. Multiple Sensory Impairment Is Associated With Increased Risk of Dementia Among Black and White Older Adults. J Gerontol A Biol Sci Med Sci. 2019;74(6):890-896. doi:10.1093/gerona/gly264. PMID: 30452551.

**S22*** Rexach JE, Polioudakis D, Yin A, Swarup V, Chang TS, Nguyen T, Sarkar A, Chen L, Huang J, Lin LC, Seeley W, Trojanowski JQ, Malhotra D, Geschwind DH. Tau pathology drives dementia risk-associated gene networks toward chronic inflammatory states and immunosuppression. Cell Rep. 2020;33(7):108398. PMID: 33207193.

**S23*–S26*: Atypical AD Variants (Table B)**

**S23*** Gorno-Tempini ML, Brambati SM, Ginex V, Ogar J, Dronkers NF, Marcone A, et al. The logopenic/phonological variant of primary progressive aphasia. Neurology. 2008;71(16):1227-1234. PMID: 18633132.

**S24*** Crutch SJ, Schott JM, Rabinovici GD, Murray M, Snowden JS, van der Flier WM, et al. Consensus classification of posterior cortical atrophy. Alzheimers Dement. 2017;13(8):870-884. PMID: 28259709.

**S25*** Ossenkoppele R, Pijnenburg YAL, Perry DC, Cohn-Sheehy BI, Scheltens NME, Vogel JW, et al. The behavioural/dysexecutive variant of Alzheimer's disease: clinical, neuroimaging and pathological features. Brain. 2015;138(Pt 9):2732-2749. PMID: 26141491.

**S26*** Miller ZA, Mandelli ML, Rankin KP, Henry ML, Babiak MC, Frazier DT, et al. Handedness and language learning disability differentially distribute in progressive aphasia variants. Brain. 2013;136(Pt 11):3461-3473. PMID: 24056533.

**S27*–S29*: Post-Electrocution / Excitotoxicity (Table B)**

**S27*** Wu WL, Gong XX, Qin ZH, Wang Y. Molecular mechanisms of excitotoxicity and their relevance to the pathogenesis of neurodegenerative diseases — an update. Acta Pharmacol Sin. 2025;46(12):3129-3142. PMID: 40389567.

**S28*** Andrews CJ, Reisner AD, Cooper MA. Post electrical or lightning injury syndrome: a proposal for an American Psychiatric Association's Diagnostic and Statistical Manual formulation with implications for treatment. Neural Regen Res. 2017;12(9):1405-1412. PMID: 29089977.

**S29*** Andrews CJ, Reisner AD. Neurological and neuropsychological consequences of electrical and lightning shock: review and theories of causation. Neural Regen Res. 2017;12(5):677-686. PMID: 28616016.
